# Cortical alpha rhythms interpolate occluded motion from natural scene context

**DOI:** 10.1101/2024.10.11.617824

**Authors:** Lu-Chun Yeh, Max Bardelang, Daniel Kaiser

**Affiliations:** Neural Computation Group, Department of Mathematics and Computer Science, Physics, Geography, Justus Liebig University Gießen, Germany; Center for Mind, Brain and Behavior (CMBB), Philipps University Marburg, Justus Liebig University Gießen, and Technical University Darmstadt, Germany

**Keywords:** Object permanence, biological motion perception, alpha oscillations, multivariate pattern analysis, cortical feedback

## Abstract

Tracking objects as they dynamically move in and out of sight is critical for parsing our ever-changing real-world surroundings. Here, we explored how the interpolation of occluded object motion in natural scenes is mediated by top-down information flows expressed in cortical alpha rhythms. We recorded EEG while participants viewed videos of a person walking across a scene. We then used multivariate decoding on alpha-band responses to decode the direction of movement across the scene. In trials where the person was temporarily occluded, alpha dynamics interpolated the person’s predicted movement. Critically, they did so in a context-dependent manner: When the scene context required the person to stop in front of an obstacle, alpha dynamics tracked the termination of motion during occlusion. As these effects were obtained with an orthogonal task at fixation, we conclude that alpha rhythms automatically interpolate occluded motion in a context-dependent way.

## Introduction

In real life, objects keep moving in and out of occlusion (e.g., pedestrians are occluded by cars passing by). Though out of sight, occluded objects are not out of our minds: an object’s behavior during an occlusion event is routinely interpolated by human observers (1), allowing for predictions about its future behavior (2). This ability, referred to as object permanence, can already be observed in infants (3). Given how fundamental the ability to keep track of occluded objects is, relatively little is known about the underlying neural mechanisms.

Previous neuroimaging studies show that the motion trajectory of occluded objects is interpolated in early visual cortex (4,5) and sustained during the initial occlusion period (6,7). Although these studies suggest a sustained representation of motion during occlusion, they probed occlusion with highly artificial stimuli like basic shapes moving on a blank background.

The way information about occluded objects is represented is likely different in natural scenes, where context constrains object movement (e.g., people can only move in certain ways across a scene). Such context-specific interpolation effects in natural scenes should be mediated by top-down predictions in the visual system. Given that previous studies suggest a link between top-down connectivity and alpha rhythms (8–10), an intriguing hypothesis is that alpha rhythms are critically involved in the interpolation of occluded motion in natural scenes.

To test this hypothesis, we applied multivariate pattern analysis to time-frequency-resolved EEG data recorded while participants viewed videos of a person walking across a scene. Critically, in some trials, the person dynamically moved in and out of occlusion, allowing us to track the interpolation of motion information while the person was out of sight.

## Results

Participants (n=48) watched 4-second videos of a person walking across a scene (either left-to-right or right-to-left) while performing an unrelated task at fixation (Fig. 1A). The person could walk across the scene without occlusion (visible condition) or with an occluder present throughout the video, which occluded the walking person from 1.5 to 3 seconds (occluded condition). Additionally, the walking person was shown on a gray background (isolated condition). Performing multivariate decoding analysis on time- and frequency-resolved EEG response patterns, we could test how object motion during occlusion is represented in patterns of alpha power (8-12hz).

**Figure 1.**
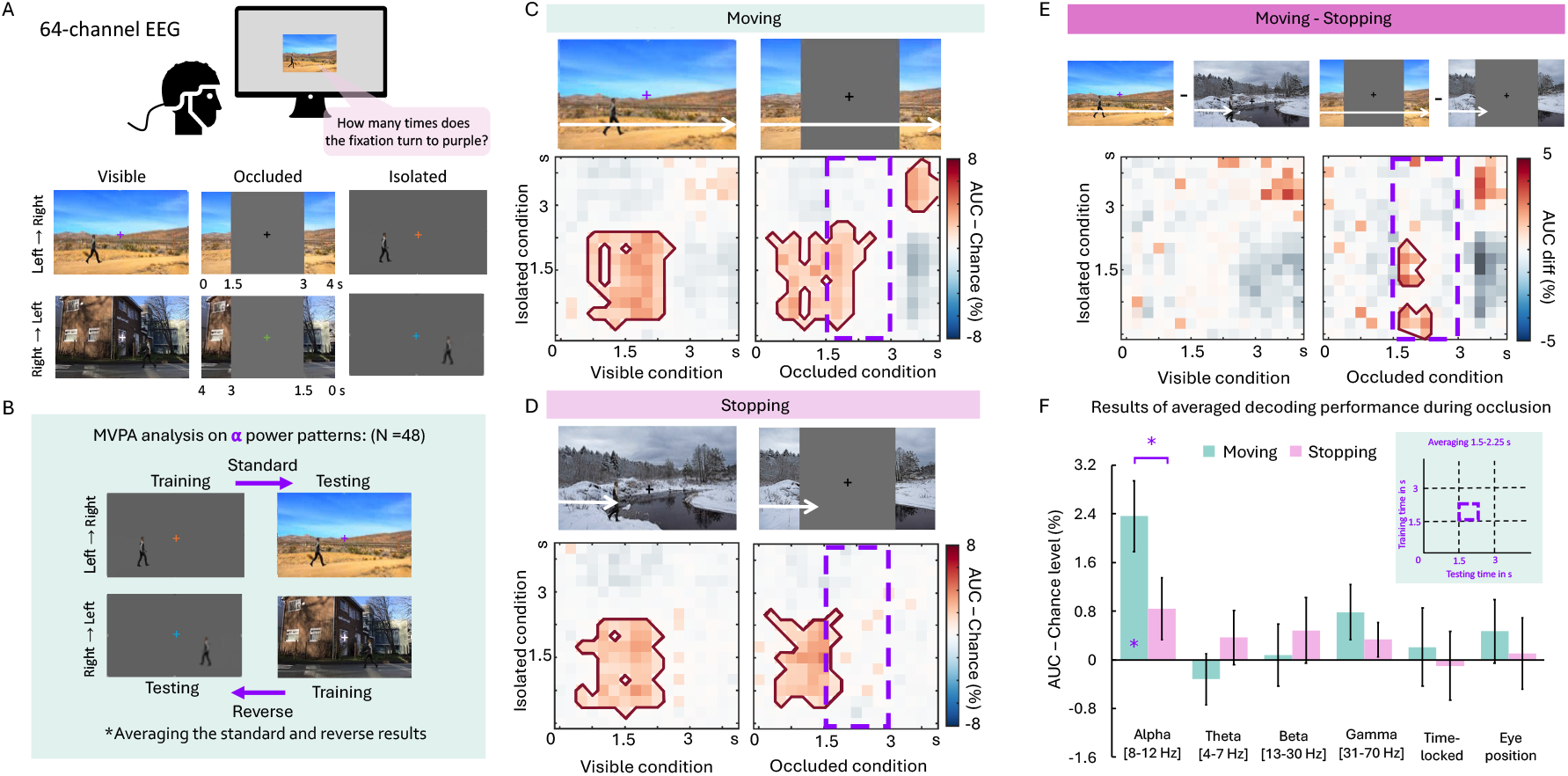
**(A)** Participants viewed 4-second videos of a person walking left-to-right or right-to-left across a scene while performing an unrelated task at fixation. In half of these trials, the person was visible throughout, while in the other half, it was occluded between 1.5s and 3s. In separate blocks, the person walked across an isolated background, used as a scene-independent benchmark for classification analysis. **(B)** We trained and tested linear classifiers on alpha-power patterns to discriminate rightward- and leftward-walking. Classifiers were trained on the isolated condition and tested on the scene conditions, and vice versa. **(C)** In the moving condition, where the person walked uninterruptedly across the scene, alpha rhythms represented motion direction in both the occluded and non-occluded conditions. The purple dashed frames indicate the occluded period, dark red lines indicate significant decoding (*p* < 0.05, corrected). **(D)** In the stopping condition, where the person stopped in front of a natural obstacle, alpha rhythms tracked the termination of movement although it happened behind the occluder. **(E)** In the occluded trials, the representation of motion was more sustained in the moving than in the stopping condition. Dark red lined indicate significant decoding differences (*p* < 0.05, corrected). **(F)** Results of averaged decoding performance within the early occlusion time window revealed that only alpha activity tracked interpolated motion during occlusion. Asterisks indicate *p* < 0.05. Error bars represent SEM.

### Alpha rhythms carry stimulus-specific information during occlusion

We trained linear classifiers on time-frequency response patterns to discriminate rightward- and leftward-walking in the isolated condition and tested them on the visible and occluded conditions (and vice versa; Fig. 1B). This cross-decoding approach allowed us to isolate the representations of object motion while removing the contribution of the background.

We first performed this analysis in a “moving” condition, where the person walked across the scene in an uninterrupted way (Fig. 1C). Classifiers trained on alpha power patterns successfully discriminated walking direction in the visible condition. Critically, when the person was occluded, decoding extended into the occlusion period (i.e., from 1.5 in the occluded condition, Fig. 1C). This suggests that motion representations in alpha rhythms are dynamically sustained into an occlusion event. However, the effect observed in the occluded time window might simply reflect a carryover of earlier representations of motion when the object was still visible. We address this concern in the following analysis.

### Alpha dynamics reflect the inferred behavior of occluded objects

To test whether decoding during occlusion was indeed related to an interpolation of motion information, we introduced an additional “stopping” condition where the person stopped in front of a natural obstacle in the scene (e.g., a river; Fig. 1D). If the brain utilized this contextual information, motion information should not be sustained during occlusion, because the person is required to stop right after disappearing behind the occluder. As expected, alpha dynamics tracked the termination of motion in this condition (Fig. 1D). The stopping condition yielded significantly lower decoding during occlusion than the moving condition (Fig. 1E). These results suggest that representations during occlusion are not driven by residual representations of the visible object. Instead, our brain actively interpolates the behavior of the included person given the current scene context.

### Only alpha rhythms interpolate occluded information

Critically, motion representations during occlusion were absent in theta, beta, and gamma rhythms, as well as in time-locked broadband responses: When comparing averaged decoding performance during the early occlusion time window (i.e., the first half of the occlusion period; Fig. 1F), only alpha rhythms yielded a significant representation of occluded motion in the moving condition (*t*(47) = 4.06, FDR corrected *p* = .001), as well as a significant difference between the moving and stopping conditions (*t*(47) = 2.73, FDR corrected *p* = .027). For detailed results across all frequency bands, see SI appendix. Furthermore, no significant effects were found when decoding was performed on participants’ eye positions recorded during the occlusion time window (Fig. 1F), suggesting that the interpolation effects in alpha rhythms were not related to differences in gaze during occlusion.

## Discussion

Our findings show that the visual system generates representations of occluded motion based on object behaviors inferred from the current scene context. This interpolated motion information is specifically represented in alpha rhythms, suggesting that alpha routes contextual motion predictions upstream (8–10).

Scene context effectively constrains the interpolation of object motion. In the stopping condition, context dictated that the walking person had to stop in front of an obstacle. In occlusion trials, the termination of motion happened after the person disappeared behind the occluder, so that the stopping had to be entirely inferred from context. Alpha dynamics tracked this inferred termination of motion, suggesting that top-down predictions accurately track the likely behavior of objects in the current environment – as opposed to a simple interpolation of a linear motion trajectory. This even held true in the absence of a task that requires interpolation.

Alpha rhythms, however, did not exhibit a difference between the moving stopping difference conditions without occlusion. One possible explanation is that in the visible condition, the person still faced the walking direction, and thus continued to share some visual features with the isolated condition. Alternatively, alpha may specifically interpolate motion based on context when feedforward signals are unavailable during occlusion, while a continued representation of motion is not needed when the whole scene is constantly present.

Consistent with previous MEG results obtained with simple visual stimuli (6), interpolated motion was not represented throughout the whole occlusion period. This may be due to interpolation becoming less accurate over time or due to representations (and consequently EEG scalp patterns) prominently changing as the person crosses the vertical midline and representations shift across hemispheres.

The involvement of alpha in interpolating occluded motion bears a resemblance to other key functions of visual alpha rhythms. First, alpha has been linked to the deployment of spatial attention across the visual field (11). The deployment of covert spatial attention may indeed play a prominent role in tracking objects during occlusion. However, the relatively broad temporal generalization pattern observed here may point towards representations of motion direction rather than reorientations of attended locations (which should result in narrow temporal generalization pattern). Second, alpha rhythms have been linked to mental imagery, where features of imagined visual contents (e.g., objects and scenes) can be retrieved from alpha activity (12,13). To which extent attention- and imagery-related mechanisms play a prominent role in the interpolation of occluded information needs to be explored in future work.

While we demonstrate that alpha rhythms carry information about the movement of occluded objects in scenes, some interesting questions remain open. First, we still need to better understand how this information is routed across visual cortex. Are alpha rhythms carrying detailed information about object features to early visual cortex, as suggested by fMRI work on occlusion (14,15)? Second, we only investigated the interpolation of object motion. Are other features of occluded objects (e.g., color, shape, or category) also represented in alpha rhythms?

Together, our study reveals alpha rhythms actively interpolate missing information in natural scenes in a context-specific way. This supports the assumption that interpolation relies on cortical top-down predictions carried by alpha rhythms.

## Materials and Methods

### Participants

Fifty-two volunteers (32 females, mean age = 26.23, SD = 4.60 years) participated in the EEG experiment. All participants had normal or corrected-to-normal vision. The participants gave informed written consent and received 10 euros per hour for their time. The study was approved by the Ethics Committee of the Julius Liebig University Gießen and was in accordance with the 6th Declaration of Helsinki. Four participants were excluded due to the low behavioral accuracy or prominent horizontal eye movements during the task, as indicated by the eye-tracking data.

### Stimuli

Participants passively viewed 4-second videos (428 × 240 pixels, 11.6*6.6° visual angle, frame rate 30 fps), split into four conditions: (1) a person walking across a scene, (2) a person walking across a scene with an occluder present, (3) a person walking across a scene but stopping in front of a natural obstacle located near the center of the scene, and (4) a person walking across a scene but stopping in front of a natural obstacle located near the center of the scene with an occluder present. In separate blocks, we showed a person walking across a blank background. For all conditions, in half of the trials, the person walked from left to right, and in the other half of trials, they walked from right to left.

The videos were created by combining eight scenes from four categories (street, grassland, snowfield, and desert, each with or without an obstacle; taken from Google Images) with a walking-person animation. The same animation was used for all scenes. The videos were initially created with a leftward walking direction and were horizontally flipped to create versions with a rightward walking direction. This yielded a total of 16 unique videos. These videos were divided into four stimulus sets, which each featured 4 videos. In these 4 videos, the leftward and rightward motion and the presence of an obstacle were uniquely associated with a specific scene category (e.g., in the street scene, the person moved from left to right, with an obstacle present). Each participant only saw one of the stimulus sets. This was done so that the behavior of the walking person could be inferred from the scene context (e.g., on the street scene, participants could predict that the person had to stop due to the obstacle).

Critically, half of the scene videos were presented with a gray occluder, which occluded the central half of the scene (and occluded the walking person between 1.5 and 3 seconds after stimulus onset). The occluder remained on the video for the whole presentation duration.

### Paradigm

Participants comfortably sat in an illuminated room at a distance of 57 cm from the monitor. EEG and eye movement data were recorded while participants performed a fixation task unrelated to the videos, counting how many times the fixation turned purple. The fixation color started at white and changed every 500ms during the video, that is, seven times during each video. It changed to purple between 1 and 4 times each trial. After the videos, a question and two numbers were presented on the screen, and participants had to press the button “F” (mapping to the number on the left side) or “J” (mapping to the number on the right side) to choose their answers.

Prior to the experiment, participants performed a calibration and validation routine to set up the eye tracking. Participants were instructed to maintain central fixation in the subsequent experiment. Next, participants practiced for two blocks of eight trials (four trials for the visible condition, followed by four trials for the occluded condition). This practice allowed participants to learn the association between scene categories and the behavior of the walking (walking uninterruptedly vs. stopping in front of an obstacle).

During the experiment, participants first performed six blocks of 32 trials with the person walking across a scene. Each block featured 16 trials for the visible and 16 trials for the occluded condition, as well as 16 trials for the moving and 16 trials for the stopping condition. Trials within a block were presented in a random order. Afterward, participants practiced four trials and performed four blocks of 32 trials for the isolated condition.

Each trial started with a fixation cross presented at the center of the screen for 500 ms. Then, a four-second video with a fixation changing its color every 500 ms was presented, followed by a response screen with the question and two numbers on the right and left. Participants responded by pressing the F key to choose the number on the left or the J key to choose the number on the right. The intertrial intervals were randomly among 800, 1,000, and 1,200 ms. The full experiment lasted around 70 minutes.

### EEG data acquisition and preprocessing

Electrophysiological data were recorded using an Easycap system with 64 channels and a Brain Products amplifier with 1000 Hz sampling rate. The electrodes included seven sites in the central line (Fz, FCz, Cz, CPz, Pz, POz, and Oz) and 28 sites over the left and right hemispheres (FP1/FP2, AF3/AF4, AF7/AF8, F1/F2, F3/F4, F5/F6, F7/F8, FC1/FC2, FC3/FC4, FT5/FT6, FT7/FT8, FT9/FT10, C1/C2, C3/C4, C5/C6, T7/T8, CP1/CP2, CP3/CP4, CP5/CP6, TP7/TP8, P1/P2, P3/P4, P5/P6, P7/P8, PO3/PO4, PO7/PO8, PO9/PO10, and O1/O2). AFz served as the ground electrode, and Fz served as a reference electrode. FP1/2 and FT9/10 were re-mounted to vertical and horizontal eye movement recorders and removed from the subsequent EEG analysis. Impedance of all the electrodes was kept below 20 kΩ. Triggers were sent from the presentation computer to the EEG computer via a parallel port.

EEG data were preprocessed offline using the Fieldtrip toolbox (16) in MATLAB (MathWorks). First, noisy channels (1.63 ± 1.01 channels) were removed by visual inspection and repaired by the mean signals of the neighboring channels. Then, artifacts caused by eye movements and blinks were removed using an independent component analysis (ICA) and visual inspection of the resulting components from the data of each participant. After that, EEG data were re-referenced using the average of all electrodes (except EOG, FP1, FP2, FT9, and FT10) and epoched from -0.5 to 4.5 s relative to stimulus onset. Epochs were baseline-corrected from -100 to 0 ms.

### EEG time-frequency analysis

Time-frequency analysis was performed using the Fieldtrip toolbox. Continuous Morlet wavelet transformation with a 7-cycle length was used for time-frequency decomposition from 0 to 4 seconds in 50-ms steps. Power values in each frequency band (4 - 7 Hz for the theta band, 8 -12 Hz for the alpha band, 13 - 30 Hz for the beta band, and 31 - 70 for the gamma band) were averaged for the following decoding analysis.

### EEG decoding analysis

To track the neural representation during dynamic occlusion, we conducted a multivariate classification analysis on time-frequency-resolved EEG data using the Amsterdam Decoding and Modeling toolbox (ADAM; 17). Linear discriminant analysis (LDA) classifiers were employed to decode moving direction (right-to-left versus left-to-right). To increase the signal-to-noise-ratio, the time-frequency-resolved data was resampled to 250ms resolution with shape-preserving piecewise cubic interpolation. The classification analysis was carried out from video onset to 4 seconds after onset, with 250-ms resolution. The Area Under the Curve (AUC) served as the measure of decoding sensitivity, where a higher AUC indicates better discrimination between classes.

A cross-decoding approach was applied to isolate motion direction information from contributions of the scene background. To this end, we probed the generalization of classifiers from the person walking on a blank background to the person walking across a scene. Specifically, classifiers were trained on discriminating motion directions in videos with gray backgrounds and testing on videos with scenes, or vice versa. Results were averaged across both train/test directions.

To assess if the resulting decoding performance exceeded chance performance, we used a cluster-based nonparametric permutation test (18). The same test was used for comparing decoding performance across conditions. Averaged decoding performance (AUC – 50%) during the early occlusion time window (from 1.5 to 2.25 seconds, i.e., analysis windows centered on 1.75 and 2.0 s) was assessed using one-tailed t-tests against zero with FDR correction. Simple effect analyses were conducted to compare the moving and stopping conditions during occlusion.

### Eye Movement Analysis

To assess the fixation stability for our paradigm, the eye-tracking data were collected during the task, except for the first two participants. Based on visual inspection, we first excluded participants who showed prominent horizontal eye movements during the task from all analyses. We then calculated the mean and SD of the horizontal eye movement across trials for each timepoint (8ms resolution) and separately for each condition. The mean horizontal eye position was within ± 0.2° from the center of the screen for each condition during the entire video, indicating stable fixation along the horizontal axis. To exclude that the interpolation effects during occlusion found in EEG signals were related to differences in eye positions, we employed Linear discriminant analysis classifiers to decode movement direction (right-to-left or left-to-right) from the eye-tracking data (left and right eye’s horizontal and vertical positions) during the first half of the occlusion period. Akin to the EEG decoding analysis, we performed the decoding on eye data from two time windows of 500ms, centered on 1.75 and 2 seconds after onset, and averaged decoding performance across these two windows (see Fig. 1F).

## Acknowledgments

L-C.Y. is supported by the Marie Skłodowska-Curie Actions (MSCA) programme (Grant agreement ID: 101149060). D.K. is supported by the DFG (SFB/TRR135, project number 222641018; KA4683/5-1, project number 518483074, KA4683/6-1, project number 536053998), “The Adaptive Mind”, funded by the Excellence Program of the Hessian Ministry of Higher Education, Science, Research and Art, and an ERC Starting Grant (PEP, ERC-2022-STG 101076057). Views and opinions expressed are those of the authors only and do not necessarily reflect those of the European Union or the European Research Council. Neither the European Union nor the granting authority can be held responsible for them.

## Author Contributions

L-C.Y. and D.K. designed the research. L-C.Y. and M.B. performed the research and analyzed the data. L-C.Y. and D.K. wrote the paper.

## Competing Interest Statement

The authors declare no competing interest.

## Supplementary Results

### EEG decoding results on other frequency bands and time-locked patterns

In the visible moving condition, we found significant clusters of motion decoding in the theta and beta bands, as well as in time-locked patterns (See *SI*. Fig 1A). We observed similar patterns across conditions prior to the occlusion time window (i.e., before 1.5 s). To systematically assess different conditions in this pre-occlusion period, we averaged decoding performance during this period (from 0 to 1.5 seconds, i.e., analysis windows centered on 0.25 to 1.25 s) and tested decoding against chance. We found significant decoding for the beta band (FDR corrected *p* <.05), marginally significant decoding for the theta band (occluded condition FDR corrected *p* = 0.057), as well as for time-locked broadband responses (FDR corrected *p* <.05). There were no significant differences between the visible and occluded conditions in any of the frequency bands or in the time-locked responses (See *SI*. Fig 1B).

**SI. Fig 1.**
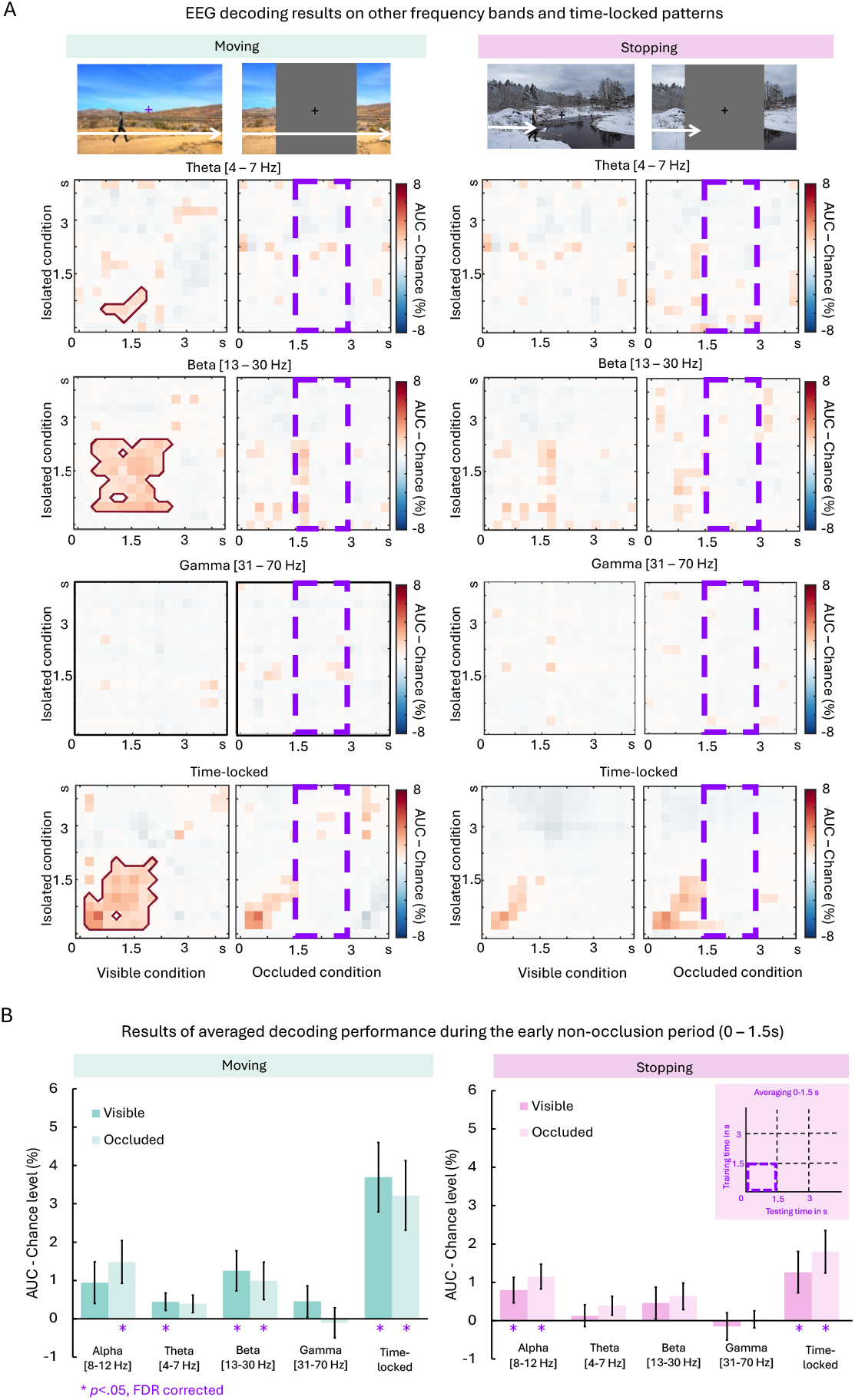
(A) Decoding results for other frequency bands and time-locked broadband patterns. We trained and tested linear classifiers on power patterns (in the alpha, theta, beta, and gamma range) or on evoked broadband responses (downsampled to 250ms resolution) to discriminate rightward- and leftward-walking. Classifiers were trained on the isolated condition and tested on the scene conditions, and vice versa. Then, we averaged the results from the two decoding directions and tested decoding against chance level using a cluster-based permutation-dependent t-test. The purple dashed frames indicate the occlusion period; dark red lines indicate significant decoding (p < 0.05, corrected). Tests for differences across conditions did not yield any significant findings. (B) Results of averaged decoding performance during the early non-occlusion period revealed walking direction can be decoded from alpha, theta, and beta, as well as time-locked patterns. There was no significant difference between visible and occluded conditions before occlusion. Asterisks indicate p < 0.05. Error bars represent SEM.

## Notes

https://osf.io/v7esy/

### Competing Interest Statement

The authors have declared no competing interest.

### Summary of Updates

changing format and adding a supplementary figure

